# Evaluating rehabilitation progress using motion features identified by machine learning

**DOI:** 10.1101/2020.04.15.043919

**Authors:** Lei Lu, Ying Tan, Marlena Klaic, Mary P. Galea, Fary Khan, Annie Oliver, Iven Mareels, Denny Oetomo, Erying Zhao

## Abstract

Evaluating progress throughout a patient’s rehabilitation episode is critical for determining effectiveness of the selected treatments and contributing to the evidence-based practice. The evaluation process is complex due to the inherent large human variations in motor recovery and the limitations of commonly used clinical measurement tools. Information recorded during a robot-assisted rehabilitation process can provide an effective means to continuously quantitatively assess movement performance and rehabilitation progress. However, selecting appropriate motion features for rehabilitation evaluation has always been challenging. This paper exploits unsupervised feature learning techniques to reduce the complexity of building the evaluation model of patients’ progress. A new feature learning technique is developed to select the most significant features from a large amount of kinematic features measured from robotics, providing clinically useful information to health practitioners with reduction of modeling complexity. A novel indicator that can reflect monotonicity and trendability is proposed to evaluate the suitability of kinematic features, which are derived from the collected data of a population of stroke patients participating in robot-aided rehabilitation. The selected kinematic features allow for human variations across a population of patients as well as over the sequence of rehabilitation sessions. The study is based on data records pertaining to 41 stroke patients using three different robot assisted exercises for upper limb rehabilitation. Consistent with the literature, the results indicate that features based on movement smoothness are the best measures among 17 kinematic features used to evaluate rehabilitation progress.

## I. Introduction

STROKE prevalence is anticipated to increase globally with Australia predicting more than one million survivors by 2050. The financial cost associated with stroke is more than 5 billions dollars annually in Australia [1]. For many stroke survivors some degree of motor function deficits are persistent. In particular, upper limb (UL) motor impairment is regarded as one of the most significant physical disabilities as it greatly reduces the quality of life [2], [3]. With the recent development of robotic technology, robot-assisted rehabilitation therapy is seen as an effective approach to improve motor function after stroke [4], [5]. In this paper the potential of the robot data recorded during the rehabilitation process to understand the progress of motor function rehabilitation is examined using machine learning techniques.

During the rehabilitation session, sensors in rehabilitation robotics can record data about the UL movement as executed by the patient in cooperation with the robot and under the guidance of clinicians. From these data it is feasible to extract various kinematic features that may be used to evaluate or characterize the quality of the UL movement and hence the progress made during rehabilitation as pointed out in [2]. Kinematic features including velocity, acceleration, and smoothness, are often used to characterize the movement quality of UL motor function. Results reported in the literature have demonstrated that some kinematic features might allow rehabilitation clinicians to evaluate rehabilitation progress [6].

In previous research [2], kinematic features were employed to represent the severity of the motor control deficits. Other authors [7] suggested to use kinematic features, including range of motion and interactive force, to predict functional motor outcomes of a rehabilitation process. In another study [8], a total of thirty-five features measured from a robotic device were suggested. The literature supports clearly that (1) kinematic features derived from sensor information recorded during the rehabilitation process can be used to assess rehabilitation progress and motion performance, (2) many kinematic features can be extracted from the recorded motion data, and it may be difficult to decide which features are truly meaningful.

It transpires that learning the most relevant features derived from motion data, is in itself a worthwhile objective way to describe the performance of UL movement. The features so learned, in some sense objectively, purely driven by the data, will assist the community to understand the rehabilitation process more completely [4]. The idea of feature learning is to identify the relevant and important information from a large scale dataset to reduce model complexity [9]. In many rehabilitation processes, it is unrealistic to expect that clinicians can have enough time to label the data [10], hence, feature learning within an unsupervised learning context is preferred to provide a solution.

In the context of rehabilitation, it is desirable that the cost function, or figure of merit function (1) can reflect the main purpose of UL motor function retraining, (2) tolerates human variations across a population of stroke survivors, (3) allows for variation over a sequence of rehabilitation exercises, so as to accommodate fatigue and concentration lapses, factors independent of the actual rehabilitation progress being made. These requirements were well accepted in the literature, see for example [11], [12].

To this end, a new figure of merit to assess kinematic features for performance evolution of motor function is developed. Firstly, the kinematic features have to possess a monotonicity property in that as the sequence of exercises progressed the kinematic feature had to equally demonstrate progress. Nevertheless, in order to be robust with respect to the inherent human variations, pure monotonicity has to be relaxed, resulting in a new concept of almost monotonicity property. By introducing a slack factor, the performance is allowed to vary within an acceptable small range to enhance the robustness. Secondly, the property of trendability is introduced to address the variation in the kinematic features over a population of stroke patients.

In summary, the main contribution of this paper is the introducing of a novel feature selecting technique that objectively identifies representative kinematic features to describe a robot-aided rehabilitation process. More precisely, a new figure of merit is proposed to assess the suitability of kinematic features, providing robustness with respect to inherent human variations across the rehabilitation process. The identified features can capture the progress of the rehabilitation process for any stroke patient. Finally, the selected representative features can be used in a scenario model to predict the overall performance of UL movements. From all feasible kinematic features it appears that those movement smoothness features provide the best insight into the motor function improvement of stroke patients.

The remainder of this paper is organized as follows. Section II describes the protocol used in this study. The relevant kinematic features are outlined in Section III, and the involved features have been described previously in literature. In Section IV, the new feature selecting criterion based on almost monotonicity and trendability is described. Section V presents the data analysis and the results of using this unsupervised feature learning technique. Section VI concludes the paper.

## II. Protocol

The Hand Hub study [13] prospectively evaluated the effectiveness of emerging technologies for UL rehabilitation in a population primarily comprised of stroke survivors. This study recruited 92 patients from a single centre in a metropolitan Melbourne location. The selection criteria were 1) aged over 18 years; 2) impaired UL following a neurological event such as stroke and multiple sclerosis; 3) assessed by a rehabilitation physician for UL impairments and potential benefits o f the program; 4) adequate cognition and communication skills to be able to follow the therapists instructions. The intervention consisted of participation in gaming and/or robotic technologies depending on the patient’s level of UL impairment. Those with more significant UL impairment would typically use the robotic device whilst those with greater levels of movement would use gaming technologies.

This study was a sub-group analysis of the 41 participants who used the robotic device in the Hand Hub study. The participants had a broad age distribution, with the minimum age 23 years and the maximum age 95 years. In the literature [14], [15], stroke survivors aged below 45 years were regarded as young adults, and those older than 75 years were considered as older adults. The total of 41 subjects were divided into three groups, young-age group (≤45 years, *n* = 8), middle-age group (> 45 years and < 75 years, *n* = 25), and old-age group (≥ 75 years, *n* = 8). As the middle-age group had the most participants (*n* = 25), the present study will mainly focus on analyzing the data collected from the middle-age group, which consists of 15 males and 10 females. Relevant parameters of the middle-age group include the mean age 61.6 ± 9.68 years, height 1.73 ± 0.13 m, weight 84.16 ± 14.82 kg, body mass index (BMI) 27.93 ± 3.24 kg/m^2^.

The particular rehabilitation strategy for all patients was robot-assisted using an exoskeleton (Armeo Power, Hocoma AG, Switzerland). As shown in Figure 1 (a), the Armeo Power provided a virtual reality, game playing environment, where numerous task reaching exercises can be programmed for execution by the stroke patient. These games can be set up in such a way that different aspects of the upper limb motion can be enhanced during the game playing. The experimental protocol was approved by the Human Research Ethics Committee of Melbourne Health (#2013.144).

**Fig. 1:**
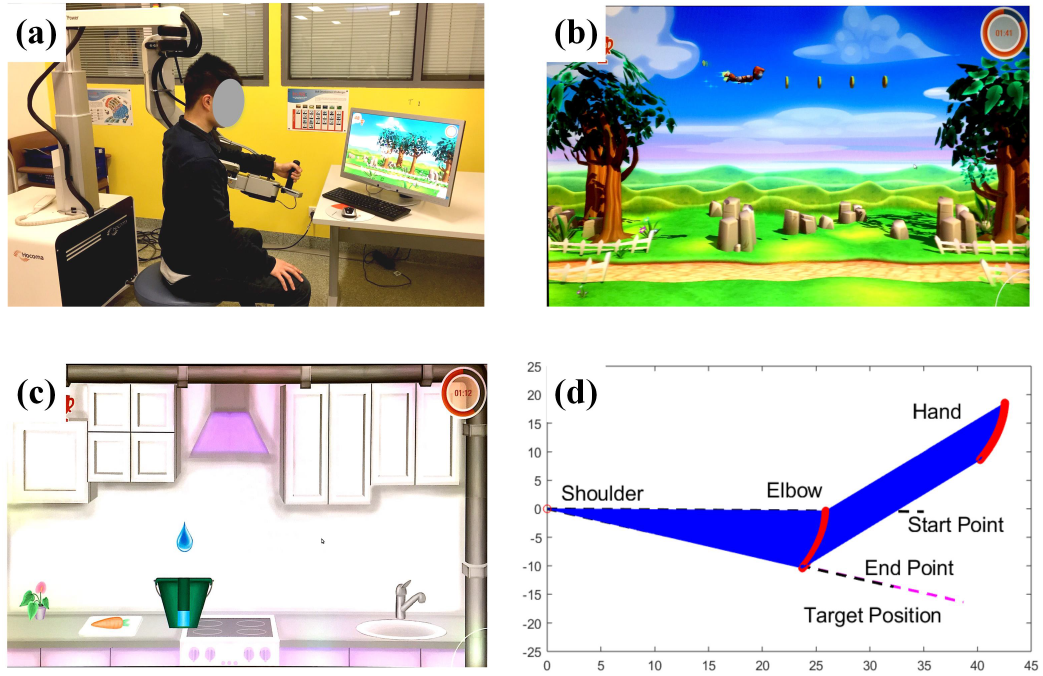
Experiment setup, (a) The Armeo Power system, (b) the EFES and EFEE games, (c) the RMS game, (d) movement trajectory.

The patients had access to several game categories, and they were working with clinicians to select which game to execute, when and for how long. As a consequence, there was great variability across the patients in terms of when, how long and which games were accessed. To select data to use as a basis for learning the relevant kinetic features, the following criteria were applied, (1) only game data of patients who used a specific game on at least 5 different dates (see also [8] for further motivation); (2) for a game that was repeatedly played in the same day, only the averaged data (over the number of games played in the day) was retained.

Using these criteria, Table I lists three different games and the corresponding patients from whom the data are used for feature learning. The date distribution and numbers of task reaching for each patient on different games involved in this study can be found in Figure 2. Illustrations of the valid games are displayed in Figures 1 (b) and (c). Figure 1 (b) presents two reaching exercises, the Elbow Flexion Extension Shoulder (EFES) and the Elbow Flexion Extension Elbow (EFEE). In the EFES game, the target is to extend and flex the shoulder, and in the EFEE game the target is to extend and flex the elbow. Both the two games require a vertical motion direction. Figure 1 (c) shows the RainMug Shoulder game (RMS) where the target is to both adduct and abduct the shoulder in a horizontal motion.

**TABLE I:** Information of valid games for the task reaching exercises

| No. | Game name | Target position | Movement direction | Index of valid participants | Total number of subjects |
| --- | --- | --- | --- | --- | --- |
| 1 | Elbow Flexion Extension Shoulder (EFES) | Shoulder | Vertical | #9, #11, #14, #15, #17, #21, #22, #23, #24, #32, #35, #38, #39 | 13 |
| 2 | Elbow Flexion Extension Elbow (EFEE) | Elbow | Vertical | #9, #11, #14, #15, #17, #21, #22, #23, #24, #32, #35, #37, #39 | 13 |
| 3 | RainMug Shoulder (RMS) | Shoulder | Horizontal | #9, #11, #15, #17, #22, #23, #24, #32, #35, #37, #38, #39 | 12 |

**Fig. 2:**
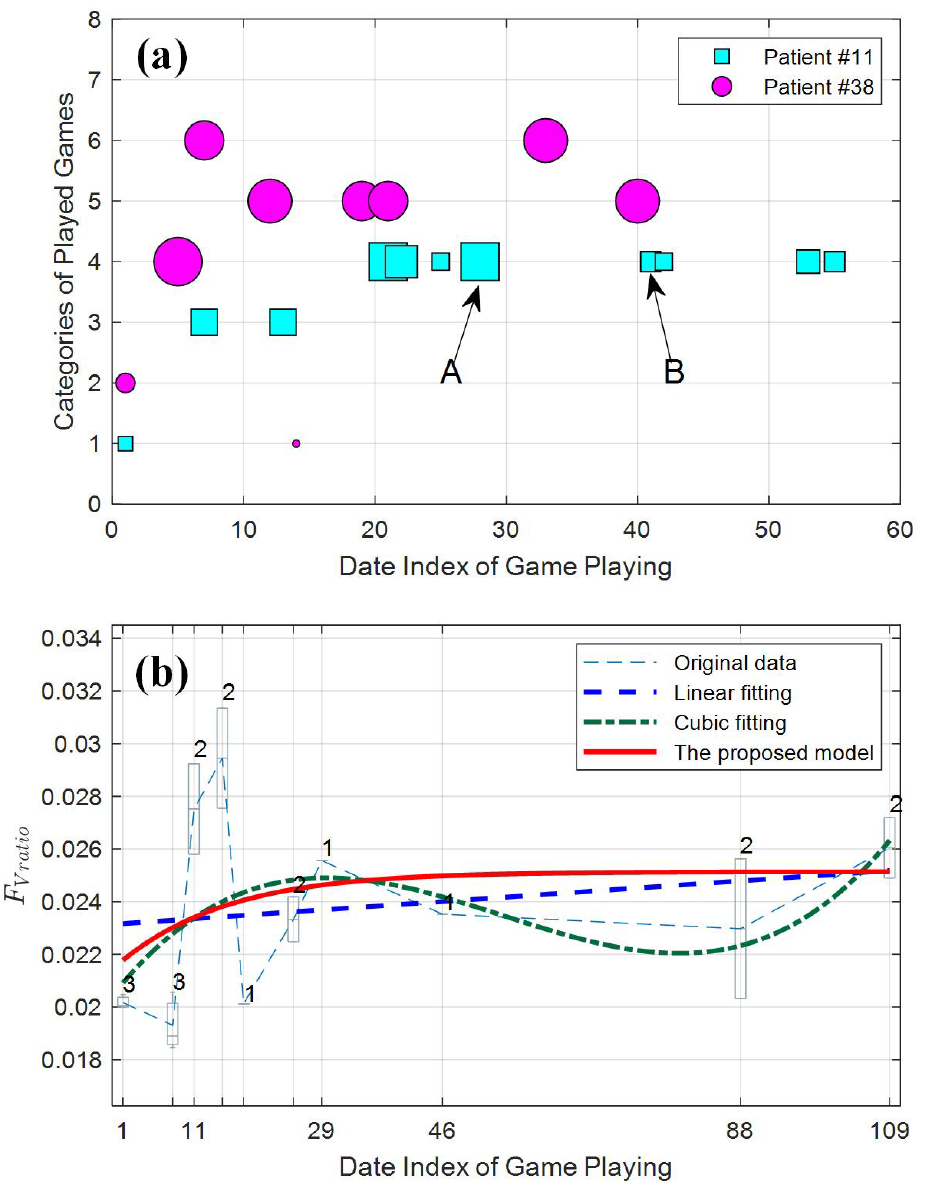
Processing of the irregularly sampled data, (a) date distribution of the task reaching exercise, (b) fitting the data.

During the motion, a total of 30 sensors in the Armeo Power system were employed to capture the UL movement, including three dimensional positions of hand, wrist and elbow, angles and torques over different joints, etc. Figure 1 (d) illustrates trajectories of hand and elbow, and the angle of shoulder flexion and extension movement for execution of one complete task reaching exercise. Next, some widely used kinematic features are calculated from the recorded data during the task reaching exercise to characterize the UL movement trajectory.

## III. Kinematic Features

Generally, clinical scores such as Fugl-Meyer Assessment, and Motor Activity Log are widely used for functional evaluation of stroke patient [3], [4]. However, these clinical measures may lack sensitivity, and require a high level of manual handling when the upper limb is densely paretic. Meanwhile, in some instances, there may be a lack of clinicians to score each particular element throughout the rehabilitation process [10]. The present study focuses on unsupervised learning to objectively identify important kinematic features from data to characterize the rehabilitation process of stroke survivors. In this section, some typical kinematic features that may be used to study UL movement are briefly described. There are many variants, and only the main ones are includeds in the text below.

### (1) Velocity and Time Features

- The maximum velocity (*F_V max_*) of the endpoint effectuator during a single execution of a movement can be seen as an indicator of motion ability [16], [17].
- Similarly, the mean velocity (*F_V mean_*) of the endpoint effectuator during motion may serve to capture progress along the rehabilitation process [16], [18].
- The time it takes to achieve the peak speed (*F_P time_*) from the rest position is expected to decrease as the rehabilitation process progresses [17], [19].
- The time it takes to execute the entire motion (*F_T dur_*) should decrease with training, and ability or ease with which the motion can be completed [17], [20].

### (2) Accuracy Features

- The global hand path ratio (*F_HP R_*) is the ratio between the length of the endpoint trajectory during the reaching movement and the minimum distance between the starting point and target [2], [21].
- The overshoot (*F_O shoot_*) is defined as the excess in the movement direction beyond the region defined by the starting point and the target [2], [21]. The smaller overshoot indicates more accurate motion control.
- Task reaching accuracy (*F_R acc_*) is defined as the residual distance between the desired target and the actual end point reached in the motion [22].
- The normalized error (*F_N err_*) is defined as the Euclidean distance between the executed position and the desired target at the end of the first sub-movement phase. This phase is defined from the starting position to the position where the speed reaches 10% of the peak speed [23].
- Normalised trajectory error (*F_Err_*) is defined as the summation of all distance differences between the desired trajectory and the patient’s trajectory. The sum is divided by the number of sample points and the minimum distance between the starting point and the target point [4].

### (3) Smoothness Features

- The number of sub-movements (*F_N sub_*) which is defined as the segment of the trajectory between successive local peaks, where these peaks are larger than 20% of the maximum speed reached during the motion [23].
- The mean jerk (*F_J mean_*) is the mean value of the third derivative of the end effector’s position [24].
- The jerk to speed ratio (*F_J ratio_*) is the ratio of the mean jerk divided by the maximum speed [16].
- Normalized jerk (*F_J norm_*) is computed by way of numerical differentiation of the trajectory’s acceleration processed by a zero phase lag low-pass filter [21].
- Spectral arc length (*F_SAL_*) is defined as the negative arc length of the amplitude and frequency-normalized Fourier magnitude spectrum of speed profile of the motion [25].
- Speed shape (*F_V ratio_*) is the ratio between the mean and maximum velocities of the endpoint trajectory. This ratio characterizes how brisk the movement is [2], [20], [24].

### (4) Force Features

- The range of grip pressure (*F_G range_*) is defined as the difference between the maximum and the minimum grip pressure observed over the motion [24].
- The skewness of the grip pressure (*F_G ske_*) is used to detect the imbalance between the subject’s ability to grip and to release the sensor [24].

## IV Unsupervised Feature Learning

### A. Notations

In this paper, the real number is denoted as ℝ, and the natural number is denoted as 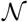. Given sensor recorded data of the task reaching exercise, the notation 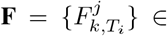 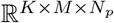 indicates the kinematic feature matrix with the element of the *k^th^* kinematic feature derived from data, which is sampled on the *T_i_* day for the *j^th^* patient, *k* = 1, 2,…, *K* is the index of kinematic features, *j* = 1, 2,…, *N*_*p*_ is index of subjects, and *T_i_* is the irregular sampling date with index *i* = 1, 2,…, *M*.

### B. Problem Formulation

Given measured movement records of stroke survivor during a rehabilitation process, a set of kinematic features can be derived from the recorded sensor information. For each participant with a session of task reaching exercise on a specific game, the kinematic feature *P_k_* can be calculated as follows,

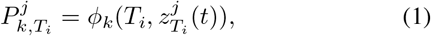

where, 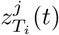 is the recorded sensor data of the session, *t* ∈ [0, *t*_*s*_] [0, *t_s_*] is the time duration of the session, *j* = 1, 2*,…, N_p_* is is the time duration of the session *j* = 1,2,… *N_p_* index of total *N_p_* subjects, *T_i_* is the irregular sampling date of the task reaching exercise, *i* = 1, 2,…, *M* is the index of total *M* sampling dates. *k* = 1, 2,…, *K* is the index of kinematic features. *ϕ*(·) is an operator that maps the timevarying signal of the session to a time-invariant feature value, such as calculating the maximum value or smoothness feature from the sensor data of the session.

Usually, the task reaching exercise consists of several individual sessions, that is, 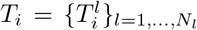, and the subject may repeat several times of the task reaching exercise. Then, an averaged kinematic feature *F_k_* of all individual sessions can be obtained as,

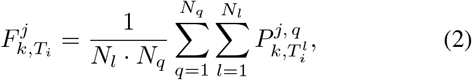

where, *l* = 1, 2,…, *N_l_* is the index of session numbers for one task reaching exercise, *q* = 1, 2,…, *N_q_* is the repeated times of the task reaching exercise during the *T_i_* day, and 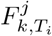 is sequence of kinematic feature that is relevant to the sampling date *T_i_* of task reaching exercise for the *j^th^* subject.

According to the research in [6], [24], the progress of motor function during rehabilitation process can be approximately indicated by kinematic features, and it is well-accepted that there are two properties of the process of stroke rehabilitation based on previous research [12], [26], [27], [28], (1) the rehabilitation process showed an average increasing trend of functional improvement over a large population, and (2) there were strong relationships of trajectories between subjects.

With these properties, we may define a suitable cost function to identify efficient kinematic features from a large datasets that contribute to the process of stroke rehabilitation. However, there usually exist large human variations and measurement uncertainties, which greatly affect the trend of the trajectory of the measured data. This paper proposed a novel concept of almost monotonicity (AM) to describe the trend of the trajectory with consideration of measurement noise and subject variations, and developed a new cost function for the unsupervised learning with combining the AM and trendability over a population of subjects. Without loss of generality, in the sequel, we denote 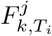 as 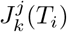 for kinematic features of interests.

### C. Suitability of a Cost Function

The suitability of a cost function for the unsupervised feature learning is a combination of AM and trendability over the population, while the AM is a relaxed version of strict monotonicity (SM). Next, the monotonicity and trendability will be introduced in details.

#### (1) Strict Monotonicity

The research in [29], [30] introduced the SM to describe the underlying trend of a cost over a population based on the assumption that

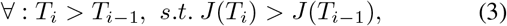

where, *T_i_* is the irregular sampling date, *i* = 2, 3,…, *M* is the date index, and *J* (*T_i_*) is the trajectory of kinematic feature.

For a single feature, without abuse of notation, we denote it as 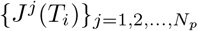 over a population of *N_p_*, the SM over this population can be calculated as following. First, we need to separate the datasets with positive increasing and negative decreasing points in the trajectory,

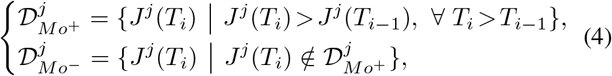

the number of data points in each of the two datasets 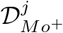 and 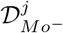 can be calculated as,

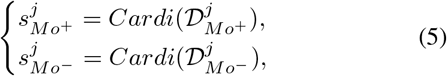

where, *Cardi*(·) is an operator to calculate the number of data points in the datasets, 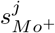 and 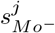 are the number of elements in the two sets. Then, the SM of the *j^th^* feature trajectory is described as,

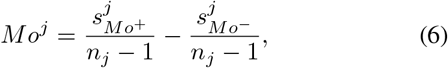

where, *j* = 1, 2,…, *N_p_* and *N_p_* is the size of the population, *n_j_* is the number of total samples (time instants) of the cost function for the *j^th^* individual in the population.

Due to large human variations and measurement noises, which usually exist in the rehabilitation process [31], the notion of the proposed AM is introduced to improve the robustness of SM.

#### (2) Almost Monotonicity

The concept of the AM is defined by relaxing the SM to a certain degree. That is, for a trajectory of kinematic feature *J* (*T_i_*), if the following condition holds,

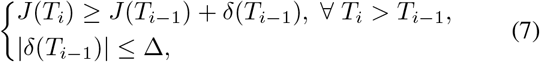

it is called almost monotonic cost function. Here, *δ*(*T*_*i* - 1_) indicates the fluctuation of the trajectory in terms of measurement noise and subject variation, and it is bounded by the term Δ, which is related to different participants and different trajectories. It is difficult to obtain the accurate value of variations due to a variety of tasks and participants, usually the variation rate can be set around 0.1, a detailed investigation of the bound is presented in Section V-D.

With the definition of AM in Eq. (7), the calculation of AM for the *j^th^* trajectory with an increasing trend, we can define the positive and negative sets accordingly,

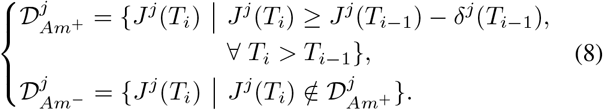

Similarly, for the trajectory *J*^*j*^(*T*_*i*_) with a decreasing trend, the positive and negative sets of the the trajectory can be calculated as follows,

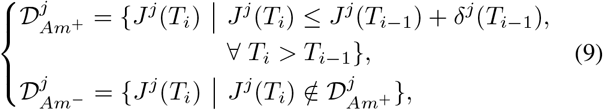

where 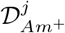 and 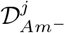 are the datasets with positive increasing and negative decreasing points in the trajectory.

Once 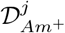 and 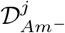 are obtained, following the similar procedure, we can obtain 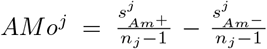, where 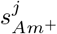 and 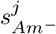 are computed similar to (5) and *n_j_* is the total number of data points in the trajectory for the *j^th^* subject.

#### (3) Trendability between Subjects

The research in [6], [32] indicates that the motor function improvement in rehabilitation process roughly follows a similar trend over a population. That is,

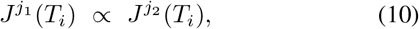

where, *j*_1_, *j*_2_ = 1, 2,…, *N_p_* and *j*_1_ ≠ *j*_2_ are subject indices, *N_p_* is the total number of subjects. The trendability is defined as the degree of progress tendency of a group of stroke survivors. The progress of each subject is characterized by the trajectory of the cost. For the 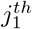 subject and the 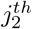 subject the trendability between them can be calculated as,

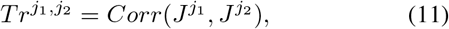

where, *Corr*(·,·) is an operator to calculate the correlation coefficient between the trajectories from two subjects. The definition of such a correlation is given as,

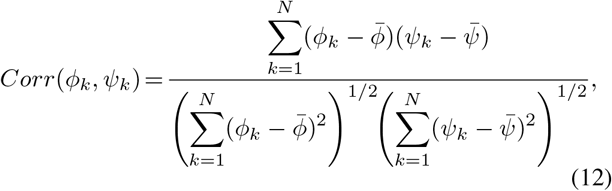

where 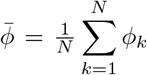 and 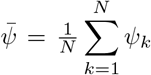 are means of two sequences respectively.

#### (4) Suitability of the Kinematic Features

By combining the AM and trendability of a cost, a new index is defined as follows to evaluate the relationship between subjects in terms of the *k^th^* kinematic feature,

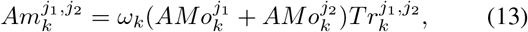

and the suitability of this cost is defined as an average over the population,

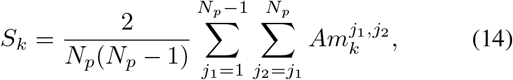

where, *S_k_* is the suitability for the *k^th^* kinematic feature, 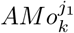 and 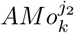 are the calculated AM values for the 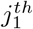 and 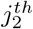 subject, respectively. 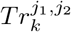 is the similarity between two two participants. *ω_k_* ∈ {−1,0,1} is a weight factor, the value of *ω_k_* is 1 when the trajectory is an increasing trend, 1 for a decreasing trend, and 0 indicates constant.

As indicated in Eq. (14), a larger AM value and a stronger trendability between costs of trajectories indicate a higher suitability value. Then, the task of selecting *n* representative features from the high dimensional data set 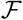 can be regarded as the following optimization problem,

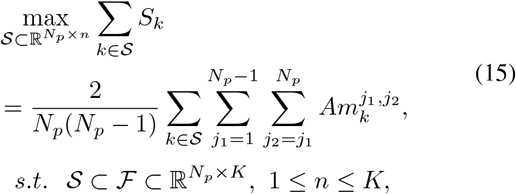

where, *S_k_* is the suitability value of the *k^th^* kinematic feature, 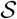 is the identified datasets with *n* representative features, 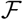 is the original datasets with total *K* kinematic features.

## V Analysis of the Collected Data

### A. Data Preprocessing

Data preprocessing was used to process the collected data during each movement session of the task reaching exercise. The data was filtered by a low-pass Butterworth filter with the cut-off frequency of 10 Hz to remove noise artefacts [17], [21]. Outliers in the measurements were identified by the three standard deviations rule and such measurements were replaced with the nearest, non-outlier point. After data preprocesing, the kinematic features detailed in Section III were calculated from the trajectory of each session of the task reaching exercise for the evaluation study.

### B. Fitting Irregularly Sampled Data

As discussed Section II, due to many factors exist in the rehabilitation process, such as the availability of patients, clinicians, and the device, it is usually difficult to sample the patient’s data in a regular way. Figure 2(a) shows the date distributions of patients #11 and #38, the *x*-axis indicates of game playing within the same day. As indicated by point *A*, patient #11 played a total of 4 different games with 10 repetitions on the 28^*th*^ day, while as shown by point *B*, the patient played the games again on the 41^*st*^ day, but there was a gap of 13 days between the two sampling dates, which clearly indicates the irregular sampling of the data.

The trajectory of the speed shape feature (*F_V ratio_*) derived from the irregularly sampled data of the EFES game is shown in Figure 2(b). The boxplot indicates the mean value of the feature, and the associated number is the repetitions of the game in the same day. To fit the irregularly sampled data, the widely used linear fitting and cubic fitting were employed for illustration. Figure 2(b) indicates the fitted trajectories by the two methods. While as suggested by the research in [26], [27], [28], the average rehabilitation process may consist of two stages, (1) the transit stage with an improvement increasing rate for a certain period, the upper limb’s functional ability may be improved in this stage; (2) the steady stage with performance fluctuation, the functional ability may be stable or have small changes. It can be clearly seen that the two fitting methods cannot fully capture the characteristics of the two stages of the rehabilitation process, thus, a new model is needed to be developed to fit the data.

In order to characterize these principles, a parameter *ρ* is introduced to describe the rate of improvement over time by using an exponential function, which is able to satisfy the discussed characteristics of the improvement of the patient during the rehabilitation process. For the time series data of kinematic feature *J* (*T_i_*), considering a gradually increasing trend, by introducing the ratio parameter, the trajectory can be described as follows,

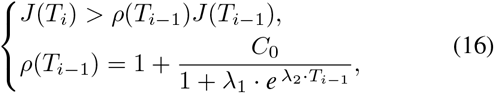

where, *ρ*(*T_i_*) is a ratio parameter to control the rate of the trajectory, *T_i_* is the sampling date of the trajectory, *i* = 1,2,…,*M*. The parameter *C*_0_ ∈ {−1, 1} is a constant value with 1 to indicate an increasing trend, and 1 for a decreasing trend. The two parameters *λ*_1_ and *λ*_2_ are used to regulate the ratio of the trajectory, and they can be obtained by optimizing the fitting process.

By using the least squares method to fit the collected data, the parameters in Eq. (16) can be identified by minimizing the regression error. The data fitting results obtained by the proposed fitting model and the other two fitting techniques are compared in Figure 2 (b), it can be clearly seen that the fitted curve by the proposed model has a gradually increasing stage in the first 29 days, and then follows a steady stage. The curve well matches the trend of the collected data, which indicates the proposed model has an excellent ability in representing the underlying trend of the irregularly sampled data.

### C. Compensation from the Robot Support

It is common that there are different levels of gravity compensation from the robot to support the rehabilitation process, which may vary among participants and the dates of game playing. Here, a weight factor is employed normalize the calculated features based on the robot support rates during the task reaching exercise. Generally, for the *k^th^* kinematic feature derived from the *j^th^* subject on the *T_i_* game playing date, the normalized feature can be described as,

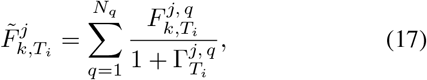

where, 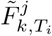 is the weighted feature for the *j^th^* subject in the *T_i_* day, and 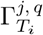 is the robot support rate for the *q^th^* game playing in the *T_i_* day, *q* = 1, 2,…, *N_q_* is the index of game playing repeated in the same day. The value of the support rate can be found in the recording of the Armeo Power system.

Take the collected sensor data from the subject #14 as an example, as shown in Figure 2 (b), the system recorded 10 separate days for the participant’s playing the EFES game and repeating the game for several times in specific days, e.g. the EFES game was played three times in the 1^*st*^ day. During each time of the game playing, the participant received compensation from the robot to support the task reaching exercise. As demonstrated by the support rates in Table II, the participant used a larger supporting rate 1.1 for the three times game playing in the 1^*st*^ and the 8^*th*^ days, while employing lower supporting rates for the other days. Then, the Eq. (17) is used to normalize all the sampled data of different days to the same level by weighting according to the supporting rate.

**TABLE II:** Supporting rate during task reaching exercise

|  |  |  |  |  |  |  |  |  |  |  |  |
| --- | --- | --- | --- | --- | --- | --- | --- | --- | --- | --- | --- |
| Date Index | 1<sup>st</sup> |  | 8<sup>th</sup> |  | 11<sup>th</sup> |  | 15<sup>th</sup> |  |  |  |  |
| Supporting Rate | 1.1, 1.1, 1.1 |  | 1.1, 1.1, 1.1 |  | 0.9, 0.9 |  | 0.8, 0.8 |  |  |  |  |
| Date Index | 18<sup>th</sup> |  | 25<sup>th</sup> |  | 29<sup>th</sup> |  | 46<sup>th</sup> |  | 88<sup>th</sup> |  | 109<sup>th</sup> |
| Supporting Rate | 0.9 |  | 0.9, 0.9 |  | 0.9 |  | 0.9 |  | 0.9, 0.9 |  | 0.9, 0.9 |

The comparison of the original data and weighted data for subject #14 is shown in Figure 3. The proposed fitting model in Section V-B is used to fit both the weighted and unweighted data, and the linear and cubic fitting are used for comparison. It should be noted that as the weighting term in Eq. (17) is larger than 1, the normalized feature value is always smaller than the original value. As illustrated in Figure 3, the results obtained by the linear fitting method are used as the baseline for both of the two types of the data, and the fitting results by the weighted data are shifted to the baseline. It can be seen that though the weighted data is more compact than the original unweighted data, the fitting curves obtained by the proposed model from the two types of data are quite same, which also indicates the model is robust against the data variation.

**Fig. 3:**
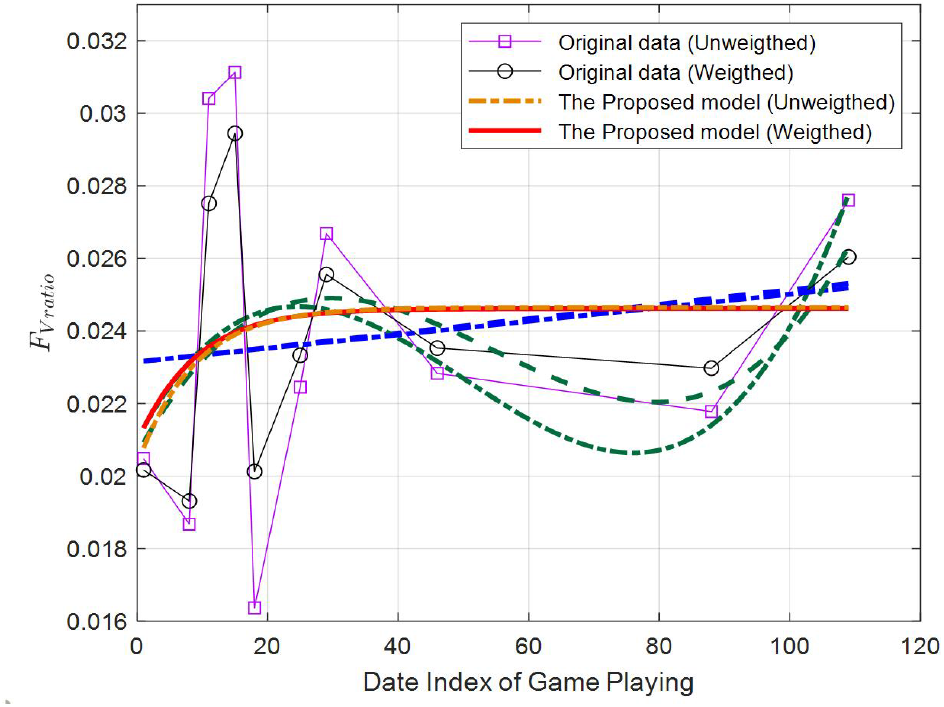
Data weighting based on the supporting rate.

After weighting the data with supporting rates, the trajectories of kinematic features for all participants can be obtained by using the proposed fitting method. Figures 4 and 5 show trajectories of the *F_V ratio_* and *F_O shoot_* features for the EFES game that were played by a total of 13 participants. The *x*-axis is the date index of game playing, and the *y*-axis indicates the calculated feature value. It can be seen that there are trajectory variations among different participants, majority of the subjects are demonstrated with increasing trends in terms of the *F_V ratio_* feature, while some of them have decreasing trends. The next section will show how to use the proposed method to evaluate the suitability of each kinematic feature.

**Fig. 4:**
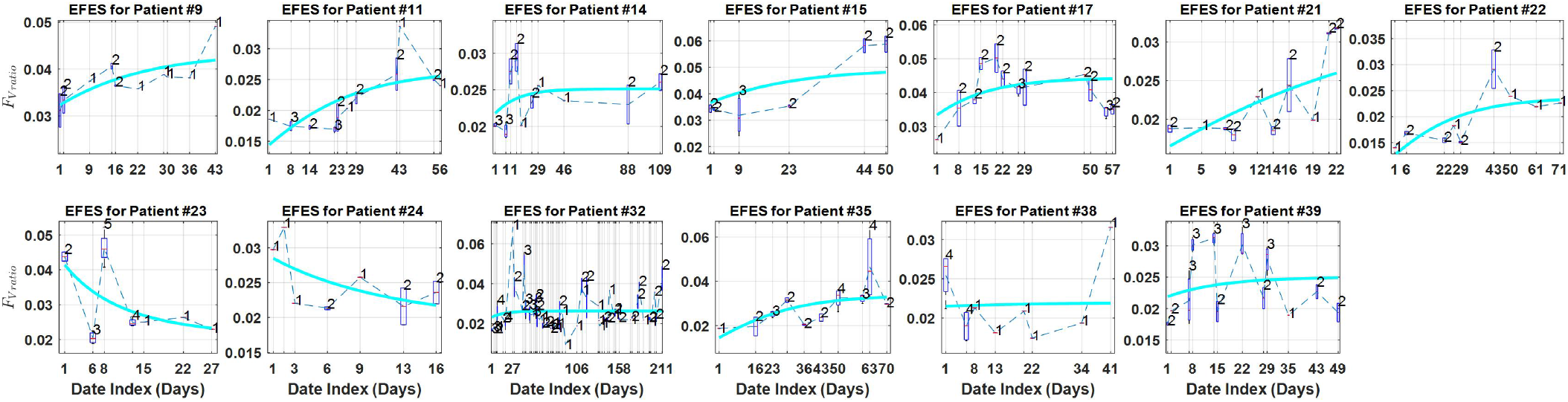
Trajectories of the *F_V ratio_* feature of the EFES game

**Fig. 5:**
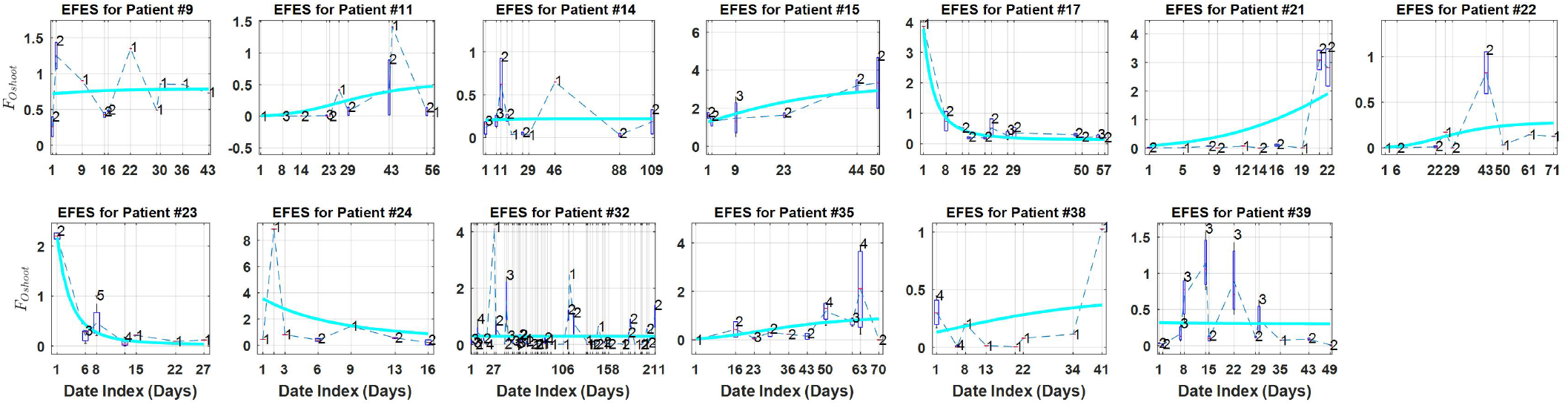
Trajectories of the *F_Oshoot_* feature of the EFES game

### D. Feature Ranking with Suitability

The proposed feature learning method is used to evaluate each kinematic feature by calculating the suitability value, which combines the AM value and similarity value of the trajectories. Table III shows the calculated AM values for the *F_V ratio_* feature of a total of 13 participants during their playing the EFES game, the relevant trajectories are shown in Figure 4. It can be seen from Table III that there are 11 positive AM values, which correspond to the increasing trends in Figure 4. There are also 2 negative AM values that are derived from the decreasing trends from participants #23 and #24. It is noted that though the trajectory of subject #38 has a decreasing trend in the first three sampling days, the overall trend of this trajectory is increasing, thus the calculated AM of the trajectory has a small positive value of 0.143, which indicates the proposed almost-monotonicity indicator is robust in capturing the underlying trend of the data.

**TABLE III:** The calculated AM value for the *F_V ratio_* feature

| Subject ID | #9 | #11 | #14 | #15 | #17 | #21 | #22 |
| --- | --- | --- | --- | --- | --- | --- | --- |
| AM Value | 0.778 | 0.8 | 0.778 | 1 | 0.455 | 0.556 | 0.5 |
| Subject ID | #23 | #24 | #32 | #35 | #38 | #39 |  |
| AM Value | -0.667 | -0.333 | 0.393 | 0.556 | 0.143 | 0.273 |  |

By calculating the AM value for each trajectory, and the correlation value between pair-wise trajectories, then the suitability value of each feature can be obtained by multiplying the two parts. Table IV shows the calculated suitability values for 10 out of the total 17 kinematic features across three games, i.e. the EFES, EFEE and RMS games. The top three maximum suitability values of kinematic features are marked with bold face. It should be noted that as discussed in Section IV-C that it is difficult to obtain the accurate value of the variation term Δ in calculating the AM value, we investigated different rates Δ_*γ*_ of the variation with 5%, 10% and 15% [33], that is, the feature is allowed a certain degree of variation of its absolute value when calculating the monotonicity increasing and decreasing points in the trajectory.

**TABLE IV:**
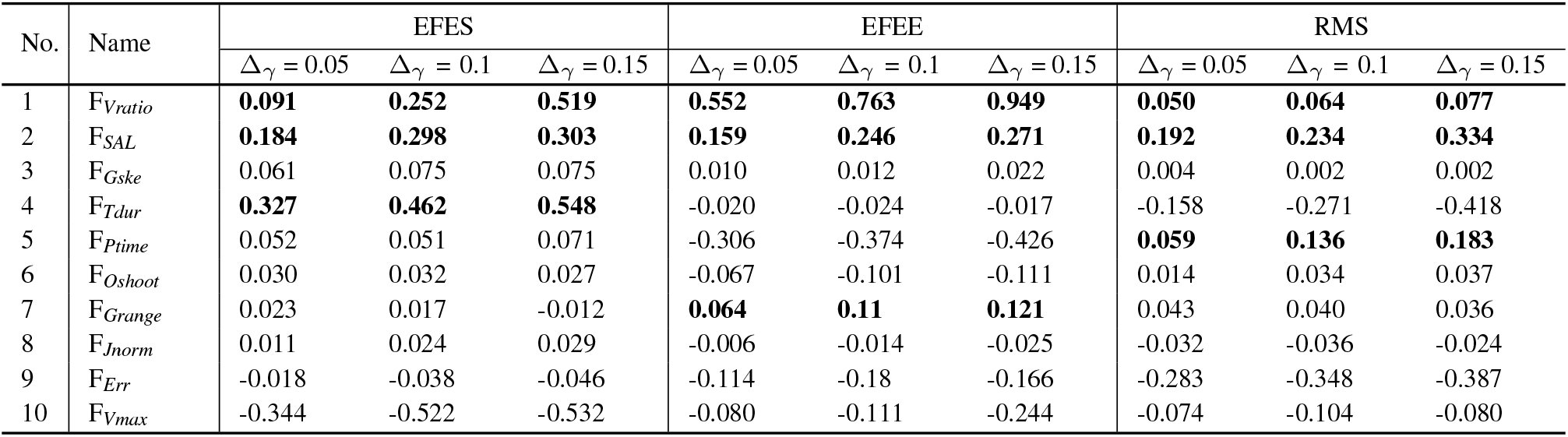
Comparison of the suitability values across three games under different variation rates

It can be seen from Table IV that with increasing the value of variation rate, the calculated suitability values for features across different games are demonstrated with increasing trends, but the category of the identified top ranking features is similar across different games. Specifically, for the EFES game, the *F_T dur_*, *F_SAL_* and *F_V ratio_* are the identified top three ranking features; for the EFEE game, the *F_V ratio_*, *F_SAL_* and *F_G range_* are the top three ranking features; and for the RMS game, the *F_SAL_*, *F_P time_* and *F_V ratio_* are the top three ranking features. It can be seen that the *F_T dur_*, *F_G range_* and *F_P time_* features are only top-ranking for a specific game, while the *F_V ratio_* and *F_SAL_* are top-ranking features and have consistent performance across all the three games, which indicates that the two feature are game-independent, and thus they are more suitable to represent the rehabilitation process.

### E. Comparison Study

The proposed method uses a novel indicator for unsupervised learning to evaluate the suitability of each kinematic feature, which combines the AM and the similarity between different trajectories. While the AM is a relaxed version of the discussed SM, meanwhile the similarity is widely used in unsupervised learning to identify representative feature [34]. To show the effectiveness, the proposed method is compared with both the SM-based and similarity-based unsupervised feature selection methods.

It should be noted that we always want to select representative features from a large database with higher suitability values. Similar to the cost definition of the AM-based method, the suitability value for the SM-based feature selection method can be defined by changing the AM term in Eq. (14) with SM,

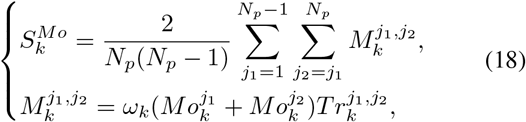

where, 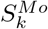 is the suitability for the *k^th^* kinematic feature, 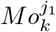 and 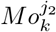 are the caluclated SM values for the 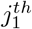 and 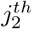 participant, respectively. *ω_k_* is a weight factor, and 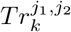 is the similarity between two trajectories. It is also used as a cost for the following similarity-based feature selection method,

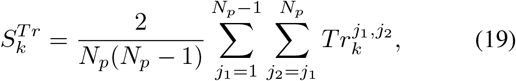

where 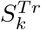 is the calculated suitability value, 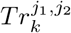 is the calculated between pair-wise trajectories as shown in Eq. (12).

With the above cost functions, the representative features can be identified with larger suitability values. Table V shows the feature evaluation results across all games by the three feature selection results. As discussed in Section V-D, the *F_SAL_* and *F_V ratio_* are the identified representative features by the proposed AM-based feature selection method. In Table V, the two features are highlighted with different colors for comparison. For our proposed AM-based method, it can be seen that the two features are identified as top three ranking features across all games. For the SM-based method, the *F_SAL_* feature has the 1^*st*^ and the 3^*rd*^ places for the three games, the *F_V ratio_* ranks the 1^*st*^ for the EFEE game, while only ranks the 4^*th*^ and 5^*th*^ for the EFES and RMS games. For the similarity-based method, it can be seen that it has the worst performance on identifying the two features, the *F_V ratio_* feature only ranks the 15^*th*^ for the RMS game. Compared with the two types of unsupervised feature selection methods, it can be clearly observed that the proposed AM-based method has excellent ability to identify top-ranking features with consistent performance across different games.

**TABLE V:**
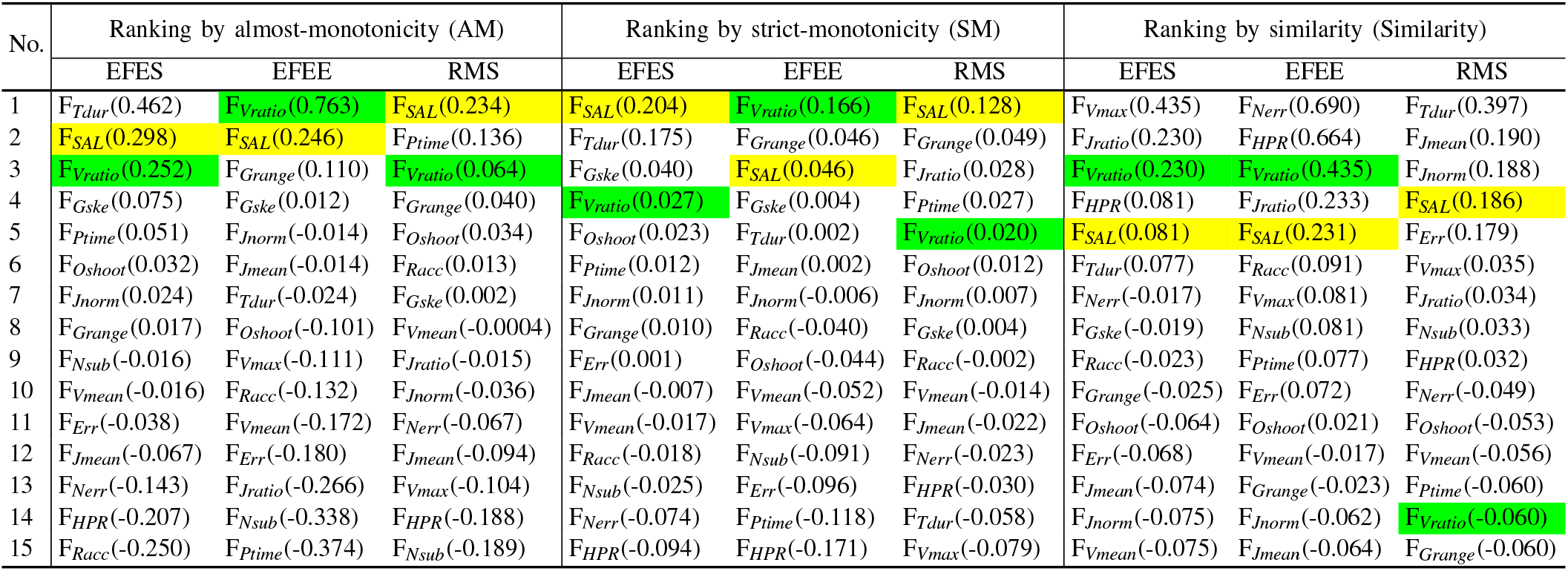
Comparison of the suitability values across different evaluation methods

### F. Discussion

With the proposed AM-based feature selection method, the identified results clearly show that the *F_SAL_* and *F_V ratio_* features are the most important features and have consistent performance in contributing to the rehabilitation process. Both the these features are regarded as smoothness indices for describing the movement performance [2], [25]. As shown in Table V, the *F_SAL_* feature ranks the 2^*nd*^ place for the EFES and EFEE games, the 1^*st*^ place for the RMS games. The *F_V ratio_* ranks the 3^*rd*^ for both the EFES and RMS games, and the 1^*st*^ for the EFEE games. It also can be seen from Figure 4 that though large human variations exist, the majority of the trajectories for the identified *F_V ratio_* feature indicate improving performance for the population of stroke patients over time, with 9 out of 13 patients demonstrating increasing trends, which indicates the robustness of the proposed feature evaluation method on identifying representative features.

It should be noted that all the listed kinematic features are widely used for analyzing the rehabilitation process in the literature, especially the accuracy features, which are usually regarded as representative features [21], [22], [23]. However, it can be seen from Table V that the accuracy features are identified with less consistent performance compared with the smoothness features. Take the *F_Oshoot_* feature for example, which is used to describe the ability of accurate motion control during movement [2], [21]. It can be seen from Table V that the *F_Oshoot_* feature ranks 6^*th*^ for the EFES game, 8^*th*^ for the EFEE game, and 5*^th^* for the RMS game. This indicates that the overshoot feature itself cannot serve as a significant contribution to the recovery during rehabilitation process. It also can been from the trajectories of the *F_O shoot_* feature for the EFES game in Figure 5 that 6 participants have increasing trends, and 3 have with decreasing trends, while the remaining 4 have very slight trends. Compared with the trajectories of the *F_V ratio_* feature as shown in Figure 4, there are 9 out of 13 participants having very clear increasing trends. Then, the calculated suitability value for the *F_V ratio_* feature is larger than that of the *F_O shoot_* feature, and thus it may be more suitable to describe the rehabilitation process.

As shown in Table V the position of the top-ranking features may have fluctuation. This is likely due to the different focuses and movement directions of the three task reaching games. The EFES and RMS game both target shoulder movements, however, the movement direction of the two games are different. The EFES game focuses on shoulder flexion/extension whilst the RMS game targets internal/external rotation. On the other hand, some features are less sensitive to different games. For example, the identified smoothness features are quite consistent for the three games. Thus, they can be used as represent features to describe the progress of a stroke patient during rehabilitation. The observations from our results are consistent with well-known results in literature [4], [6], [35], which indicated the strong correlations between smoothness indices and clinical measures, and they were useful to describe the rehabilitation process.

## VI. CONCLUSION

This paper has proposed a novel unsupervised learning method to identify key features of progress during the rehabilitation process of a group of stroke patients, with the data collected from a rehabilitation robotic device. A new indicator was developed to evaluate the suitability of kinematic features in terms of their contributions towards the progress during rehabilitation based on some standing assumptions. The proposed model was used to identify representative features from unlabeled information recorded from task reaching exercises executed by stroke patients in the presence of large human variations. The identified results indicated that smoothness indices (the spectral arc length and the speed shape features) demonstrate important aspects of recovery and they were game-independent. Moreover, the proposed method also provides a general framework of unsupervised learning for a large class of processes with some clear underlying trends.

## REFERENCES

[1] E. Appleby, S. T. Gill, L. K. Hayes, T. L. Walker, M. Walsh, and S. Kumar, “Effectiveness of telerehabilitation in the management of adults with stroke: A systematic review,” PloS one, vol. 14, no. 11, 2019.

[2] M. Longhi, A. Merlo, P. Prati, M. Giacobbi, and D. Mazzoli, “Instrumental indices for upper limb function assessment in stroke patients: a validation study,” J. Neuroeng. Rehabil., vol. 13, no. 1, p. 52, 2016.

[3] O. Einav, D. Geva, D. Yoeli, M. Kerzhner, and K.-H. Mauritz, “Development and validation of the first robotic scale for the clinical assessment of upper extremity motor impairments in stroke patients,” Top. Stroke Rehabil., vol. 18, no. sup1, pp. 587–598, 2011.

[4] O. Celik, M. K. O’Malley, C. Boake, H. S. Levin, N. Yozbatiran, and T. A. Reistetter, “Normalized movement quality measures for therapeutic robots strongly correlate with clinical motor impairment measures,” IEEE Trans. Neural Syst. Rehabil. Eng., vol. 18, no. 4, pp. 433–444, 2010.

[5] E. P. Washabaugh, J. Guo, C.-K. Chang, C. D. Remy, and C. Krishnan, “A portable passive rehabilitation robot for upper-extremity functional resistance training,” IEEE Trans. Biomed. Eng., vol. 66, no. 2, pp. 496–508, 2018.

[6] C. Bosecker, L. Dipietro, B. Volpe, and H. Igo Krebs, “Kinematic robot-based evaluation scales and clinical counterparts to measure upper limb motor performance in patients with chronic stroke,” Neurorehabil. Neural Repair., vol. 24, no. 1, pp. 62–69, 2010.

[7] R. S. Calabrò, M. Russo, A. Naro, D. Milardi, T. Balletta, A. Leo, S. Filoni, and P. Bramanti, “Who may benefit from armeo power treatment? a neurophysiological approach to predict neurorehabilitation outcomes,” PM&R, vol. 8, no. 10, pp. 971–978, 2016.

[8] H. I. Krebs, M. Krams, D. K. Agrafiotis, A. DiBernardo, J. C. Chavez, G. S. Littman, E. Yang, G. Byttebier, L. Dipietro, A. Rykman, et al., “Robotic measurement of arm movements after stroke establishes biomarkers of motor recovery,” Stroke, vol. 45, no. 1, pp. 200–204, 2014.

[9] X. Zhu, H.-I. Suk, S.-W. Lee, and D. Shen, “Subspace regularized sparse multitask learning for multiclass neurodegenerative disease identification,” IEEE Trans. Biomed. Eng., vol. 63, no. 3, pp. 607–618, 2016.

[10] E. A. Kirchner, J. C. Albiez, A. Seeland, M. Jordan, and F. Kirchner, “Towards assistive robotics for home rehabilitation,” in Biodevices, pp. 168–177, 2013.

[11] N. Nordin, S. Q. Xie, and B. Wünsche, “Assessment of movement quality in robot-assisted upper limb rehabilitation after stroke: a review,” J. Neuroeng. Rehabil., vol. 11, no. 1, p. 137, 2014.

[12] D. J. Reinkensmeyer, E. Burdet, M. Casadio, J. W. Krakauer, G. Kwakkel, C. E. Lang, S. P. Swinnen, N. S. Ward, and N. Schweighofer, “Computational neurorehabilitation: modeling plasticity and learning to predict recovery,” J. Neuroeng. Rehabil., vol. 13, no. 1, p. 42, 2016.

[13] M. Galea, F. Khan, B. Amatya, A. Elmalik, M. Klaic, and G. Abbott, “Implementation of a technology-assisted programme to intensify upper limb rehabilitation in neurologically impaired participants: A prospective study,” J. Rehabil. Med., vol. 48, no. 6, pp. 522–528, 2016.

[14] D. Leys, L. Bandu, H. Henon, C. Lucas, F. Mounier-Vehier, P. Rondepierre, and O. Godefroy, “Clinical outcome in 287 consecutive young adults (15 to 45 years) with ischemic stroke,” Neurology, vol. 59, no. 1, pp. 26–33, 2002.

[15] J. A. Falconer, B. J. Naughton, D. C. Strasser, and J. M. Sinacore, “Stroke inpatient rehabilitation: a comparison across age groups,” J. Am. Geriatr. Soc., vol. 42, no. 1, pp. 39–44, 1994.

[16] S. Straudi, E. Chew, C. Iahn, F. Fregni, and P. Bonato, “Combining transcranial direct current stimulation and gravity-supported, computer-enhanced arm training in a chronic pediatric stroke survivor: a case report,” Clin. Case Rep. Rev., vol. 2, no. 1, pp. 301–306, 2015.

[17] A. Weightman, N. Preston, M. Levesley, R. Holt, M. Mon-Williams, M. Clarke, A. J. Cozens, and B. Bhakta, “Home-based computer-assisted upper limb exercise for young children with cerebral palsy: A feasibility study investigating impact on motor control and functional outcome,” J. Rehabil. Med., vol. 43, no. 4, pp. 359–363, 2011.

[18] M. Longhi, A. Merlo, P. Prati, M. Giacobbi, and D. Mazzoli, “Instrumental indices for upper limb function assessment in stroke patients: a validation study,” J. Neuroeng. Rehabil., vol. 13, no. 1, p. 52, 2016.

[19] J. Fong, V. Crocher, Y. Tan, D. Oetomo, and I. Mareels, “Emu: A transparent 3d robotic manipulandum for upper-limb rehabilitation,” in Rehabil. Robotics (ICORR), 2017 Int. Conf. on, pp. 771–776, IEEE, 2017.

[20] H. I. Krebs, S. E. Fasoli, L. Dipietro, M. Fragala-Pinkham, R. Hughes, J. Stein, and N. Hogan, “Motor learning characterizes habilitation of children with hemiplegic cerebral palsy,” Neurorehabil. Neural Repair., vol. 26, no. 7, pp. 855–860, 2012.

[21] A. Merlo, M. Longhi, E. Giannotti, P. Prati, M. Giacobbi, E. Ruscelli, A. Mancini, M. Ottaviani, L. Montanari, and D. Mazzoli, “Upper limb evaluation with robotic exoskeleton. normative values for indices of accuracy, speed and smoothness,” NeuroRehabil., vol. 33, no. 4, pp. 523–530, 2013.

[22] J. J. Daly, N. Hogan, E. M. Perepezko, H. I. Krebs, J. M. Rogers, K. S. Goyal, M. E. Dohring, E. Fredrickson, J. Nethery, and R. L. Ruff, “Response to upper-limb robotics and functional neuromuscular stimulation following stroke,” J. Rehabil. Res. Dev., vol. 42, no. 6, pp. 723–736, 2005.

[23] A. De Luca, P. Giannoni, H. Vernetti, C. Capra, C. Lentino, G. A. Checchia, and M. Casadio, “Training the unimpaired arm improves the motion of the impaired arm and the sitting balance in chronic stroke survivors,” IEEE Trans. Neural Syst. Rehabil. Eng., vol. 25, no. 7, pp. 873–882, 2017.

[24] J. Zariffa, N. Kapadia, J. L. Kramer, P. Taylor, M. Alizadeh-Meghrazi, V. Zivanovic, U. Albisser, R. Willms, A. Townson, A. Curt, et al., “Relationship between clinical assessments of function and measurements from an upper-limb robotic rehabilitation device in cervical spinal cord injury,” IEEE Trans. Neural Syst. Rehabil. Eng., vol. 20, no. 3, pp. 341–350, 2012.

[25] S. Balasubramanian, A. Melendez-Calderon, and E. Burdet, “A robust and sensitive metric for quantifying movement smoothness,” IEEE Trans. Biomed. Eng., vol. 59, no. 8, pp. 2126–2136, 2012.

[26] V. Klamroth-Marganska, J. Blanco, K. Campen, A. Curt, V. Dietz, T. Ettlin, M. Felder, B. Fellinghauer, M. Guidali, A. Kollmar, et al., “Three-dimensional, task-specific robot therapy of the arm after stroke: a multicentre, parallel-group randomised trial,” Lancet Neurol., vol. 13, no. 2, pp. 159–166, 2014.

[27] J. van Kordelaar, E. E. van Wegen, R. H. Nijland, A. Daffertshofer, and G. Kwakkel, “Understanding adaptive motor control of the paretic upper limb early poststroke: the explicit-stroke program,” Neurorehabil. Neural Repair., vol. 27, no. 9, pp. 854–863, 2013.

[28] P. S. Lum, C. G. Burgar, P. C. Shor, M. Majmundar, and M. Van der Loos, “Robot-assisted movement training compared with conventional therapy techniques for the rehabilitation of upper-limb motor function after stroke,” Arch. Phys. Med. Rehabil., vol. 83, no. 7, pp. 952–959, 2002.

[29] J. B. Coble, “Merging data sources to predict remaining useful life–an automated method to identify prognostic parameters,” PhD Dissertation, University of Tennessee, 2010.

[30] L. Lu, Y. Tan, D. Oetomo, I. Mareels, and S. An, “Feature learning in assistive rehabilitation robotic systems,” in 2018 40th Conf. Proc. IEEE Eng. Med. Biol. Soc., pp. 2511–2514, IEEE, 2018.

[31] G. DeJong, C.-H. Hsieh, K. Putman, R. J. Smout, S. D. Horn, and W. Tian, “Physical therapy activities in stroke, knee arthroplasty, and traumatic brain injury rehabilitation: their variation, similarities, and association with functional outcomes,” Phys. Ther., vol. 91, no. 12, pp. 1826–1837, 2011.

[32] S. Prabhakaran, E. Zarahn, C. Riley, A. Speizer, J. Y. Chong, R. M. Lazar, R. S. Marshall, and J. W. Krakauer, “Inter-individual variability in the capacity for motor recovery after ischemic stroke,” Neurorehabil. Neural Repair., vol. 22, no. 1, pp. 64–71, 2008.

[33] M. R. Selwyn, Principles of experimental design for the life sciences. CRC Press, 1996.

[34] E. Hosseini-Asl, J. M. Zurada, G. Gimel’farb, and A. El-Baz, “3-d lung segmentation by incremental constrained nonnegative matrix factorization,” IEEE Trans. Biomed. Eng., vol. 63, no. 5, pp. 952–963, 2015.

[35] B. Rohrer, S. Fasoli, H. I. Krebs, R. Hughes, B. Volpe, W. R. Frontera, J. Stein, and N. Hogan, “Movement smoothness changes during stroke recovery,” J. Neurosci., vol. 22, no. 18, pp. 8297–8304, 2002.

